# Cognitive impairments in a mouse model for Huntington’s disease correlate with presymptomatic locomotion and number of CAG repeats

**DOI:** 10.64898/2026.07.17.738914

**Authors:** Olga Jung, Sabine Hoffmeister-Ullerich, Asina Omriouate, Josefine Plumhoff, Michael R. Kreutz, Katarzyna M. Grochowska, Fabio Morellini

## Abstract

Huntington’s disease (HD) is a progressive neurodegenerative disorder caused by an expanded CAG repeat in the *huntingtin* (*HTT*) gene. The disease is characterized by movement disorders, and it also presents with personality changes, including apathy and aggression, along with cognitive decline. While most animal models for HD have been validated for motor deficits, less is known about alterations in other behavioral functions. Here, we performed a longitudinal study to analyze the behavior of a knock-in mouse model of HD with a chimeric mouse/human exon 1 containing 140 CAG repeats inserted in the murine *huntingtin* gene. We specifically inquired about the onset of cognitive impairments in knock-in mice and whether changes in various behavioral functions such as locomotion, anxiety, and cognition correlate at the individual level. Our data indicate that female and male knock-in mice exhibit reductions in body weight, novelty-induced locomotion, and remote spatial memory retrieval. However, social behavior, working, and short-term memory remain unaffected. Within knock-in mice, lower open-field activity correlated with poorer remote memory performance. Moreover, CAG repeat length negatively correlated with locomotor activity and spatial memory, indicating that greater repeat expansion predicts more severe behavioral impairment. These findings identify early affective changes, followed by selective long-term memory and locomotor deficits, in knock-in mice, supporting this model as a useful platform for studying prodromal HD and repeat-length-dependent disease variability.

**Highlights:**

- CAG140 knock-in mice show early anxiety, later reduced locomotion and memory deficits
- Long-term and remote memory are impaired while short-term and working memory are spared
- Lower locomotion at the age of 8 months correlates with poorer memory at 14 months of age in individual CAG140 knock-in mice
- Greater CAG repeat length predicts worse locomotion and memory

## Introduction

Huntington’s disease (HD) is a fatal neurodegenerative disorder characterized by motor, cognitive, and psychiatric impairments. These impairments result from increased cytosine-adenosine-guanine (CAG) triplet repeats in the *huntingtin* (*HTT*) gene, located on the short arm of chromosome 4. Following translation, this mutation produces an elongated polyglutamine repeat at the N-terminus of the huntingtin protein, leading to neurotoxicity, particularly in the striatum (Snowden, 2017). At the age of onset it manifests by the appearance of typical choreiform extrapyramidal motor signs, which correlate with the number of CAG repeats: a higher number of repeats results in earlier onset. The characteristic choreiform dysfunction in HD involves abrupt, uncontrolled, and intense movements of the body and face, including dysarthrophonia and hyperlordosis. At later stages, rigidity, bradykinesia, akinesia, and decreased coordination and balance become predominant (Ghosh and Tabrizi, 2018).

Although the choreiform motor dysfunction is prominent and lends its name to the disease, slowed psychomotor function is often the earliest sign of HD (Ghosh and Tabrizi, 2018). In fact, the quality of life in HD is primarily affected by cognitive impairment. Cognitive deficits emerge years before the onset of motor symptoms and encompass a broad spectrum of domains. During the pre-symptomatic period, impairments in executive functions, such as attention, planning, organization, task-oriented strategy adaptation, and cognitive flexibility, have been reported (Giralt et al., 2012). Psychiatric symptoms, including depression and reductions in declarative, procedural, source, and prospective memory, as well as diminished recall and learning abilities, are also common. Furthermore, HD results in deficits in social cognition, including impaired recognition and expression of emotions and empathy (Adjeroud et al., 2016).

Animal models are valuable when developing and testing therapeutic tools against the disease in the preclinical stage (Upadhayay et al., 2023). Transgenic knock-in (KI) models are characterized by the replacement of the mouse *Htt* gene, in whole or in part, with the human *HTT* gene (Ramaswamy et al., 2007). They have the advantage of exploring the gradual, progressive deterioration of cognitive functions and behavior and re-creating the pathogenic progression of particular facets in humans (Döbrössy et al., 2009; Nittari et al., 2023). Among the KI models, the CAG140 KI mice were generated by inserting a chimeric *Huntington’s disease homolog* (*HDh)* gene homolog and mutated human *HTT* exon 1 into the exogenous murine *HDh* locus, under the control of the mouse *Htt* promoter (Menalled et al., 2003). From the CAG140 line emerged the zQ175 model due to a spontaneous mutation that increased the CAG repeat number: whereas the CAG140 mice have about 120-140 CAG repeats, the zQ175 mice harbor about 175-190 repeats. As predicted by the longer CAG sequence, disease onset is earlier in zQ175 mice than in CAG 140 mice, with a stronger, more robust phenotype (Menalled et al., 2012; Rising et al., 2011; Harrison et al., 2025; Rebec et al., 2022).

Despite the fact that CAG140 KI mouse is a widely used slow-progressive model of HD, the evolution of possible impairments in non-motor related behaviors associated with HD has not been comprehensively charachterized. Only a few studies have reported early hypoactivity as early as 11 weeks (Menalled et al., 2003; Fowler and Muma, 2015; Stefanko et al., 2017), increased immobility in the tail suspension and forced-swim test, and a slower reversal learning in the Y maze test (Stefanko et al., 2017). While CAG-repeat length is known to strongly predict phenotypic severity across KI cohorts (Huang et al., 2021; Alexandrov et al., 2016), it remains unclear whether natural variability in CAG-repeat length within CAG140 mice correlates with individual behavioral severity. In this study, we aim to replicate and extend previous behavioral analyses in CAG140 KI mice, focusing on memory function and examining potential cross-domain interactions among behavioral functions. In addition, we examined whether behavioral changes observed in young, symptom-free animals are associated with movement impairments in later adulthood. Our objective was to determine whether the emergence of specific symptoms predicts the onset of other behavioral impairments and memory decline, and whether these outcomes are related to CAG-repeat length at the individual mouse level.

## Materials and Methods

### 2.1 Animals and husbandry

Female and male wild-type (WT) and knock-in (KI) mice (Menalled et al., 2003) were used (Female WT= 15; Female KO= 12; Male WT= 17; Male KI= 11). Mice were transferred from a breeding facility to the experimental animal facility at 5 weeks of age and handled until they underwent their first open-field test at 8 weeks of age. The mice were housed in Type III cage groups of 4, composed of sex- and age-matched mice of the two genotypes. Five-to seven-week-old mice were handled for three consecutive days, with each mouse receiving at least 1 minute of handling per day, and body weight was measured on the first day. Body weight measurements were repeated twice during the experiment to detect changes and assess developmental differences among genotypes. The mice were tested at 8-10 weeks and 13-16 weeks, with at least 1 day between experiments (Fig. 1A). Animal experiments were approved by the local authorities of the city-state of Hamburg (Behörde für Gesundheit und Verbraucherschutz, Fachbereich Veterinärwesen; Ü001/2019, N153/2021) and the animal care committee of the University Medical Center Hamburg-Eppendorf, in compliance with German law (Tierschutzgesetz der Bundesrepublik Deutschland, TierSchG) and the guidelines of Directive 2010/63/EU.

**Figure 1.**
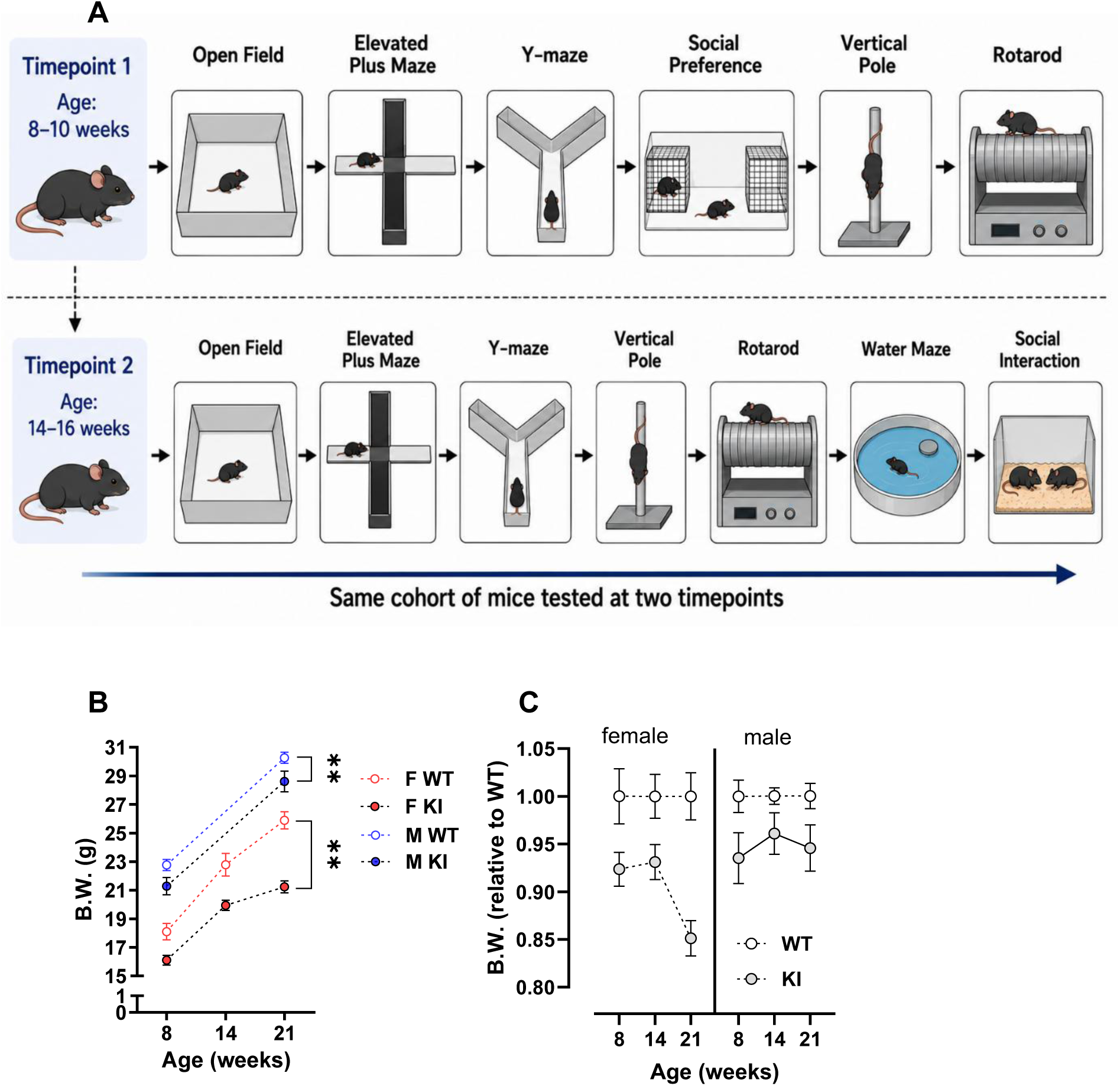
Decreased body weight in KI mice. A, experimental design of the longitudinal behavioral study. B, body weight (B.W.) measured in 8, 14, and 21-week-old mice. C, the values shown in panel (A) were normalized to the WT group within each age. **, *** p< 0.01 and 0.001, respectively (effect of genotype within each sex, mixed 2-way ANOVA).

### 2.2 Open field

The experiment was performed in a wooden box (50 x 50 x 40 cm) laminated with light grey plastic and illuminated with yellow light (25 lux). A single mouse was placed in a corner of the box and allowed to move freely for 15 minutes. Distance moved, maximal velocity, time spent in the center of the box, time spent in the border area, latency to cross the center, and distance to the wall were analyzed using EthoVision (Noldus). Rearing and self-grooming during the first 5 minutes were analyzed offline by a blinded observer using The Observer (Noldus).

### 2.3 Elevated plus maze

The plastic maze was shaped like a plus and consisted of four 30 cm-long, 5 cm-wide arms connected by a 5 x 5 cm central area. 15 cm-high walls enclosed two opposing arms, which opened to the central area, called closed arms. The other two opposing arms, the so-called open arms, were bordered by a rim 2 mm high. The maze was elevated 75 cm above the floor on a metal stand. An infrared camera was used to record the mouse’s behavior in complete darkness. The mouse was placed in the center, facing one of the open arms. The mouse’s behavior was video-recorded for five minutes and analyzed using The Observer software, as described (Brandeweide et al., 2005).

### 2.3 Spontaneous alternation in the Y-Maze

The spontaneous alternation test assesses working memory, assuming that mice are motivated to explore novel and recently unvisited environments (Buccafusco, ed., 2009). The maze consisted of three transparent Plexiglas corridors (40 cm long, 5 cm wide, and 30 cm high) forming a Y with 120-degree turns between arms. Paper sheets with different patterns were placed along each wall to help the mouse distinguish the arms. The maze was illuminated by dim light (15 lux). Mice were placed in the maze’s center and allowed to move until they completed 26 transitions between corridors after the first entry into one of the corridors or after 20 minutes.

### 2.4 Spatial object recognition

This test was designed to test short-term spatial memory. The paradigm is based on the same assumption as the spontaneous alternation test: Rodents intrinsically tend to investigate novel stimuli. The spatial object recognition test was conducted in the same box used for the open-field test, which was divided into two equal compartments by a white PVC wall with a hole in the middle, allowing the mouse to move between them. The arena was illuminated with white light (15 lux). The paradigm consisted of two trials: an exposure (10 min) and a test (5 min), with a 10 min inter-trial interval during which mice were returned to their housing cage. During the exposure trial, two identical objects, unfamiliar to the mice, were placed in one corner of each compartment. During the test trial, one of the two objects was relocated to a different corner of the compartment. Distance moved, mean velocity, and time spent at the objects were analyzed with Ethovision. Sniffing the objects, self-grooming, rearing on the wall, and rearing on objects were analyzed with The Observer.

### 2.5 Social preference

The motivation to investigate a social stimulus was assessed by giving the experimental mouse the choice between a beaker containing an unfamiliar sex-matched mouse and an empty beaker. The apparatus consisted of a square box (50 cm x 50 cm and 40 cm high) illuminated with white light (10 lux). Two beakers, made of transparent plastic (diameter 10 cm, height 15 cm), with several holes (diameter 1 cm) drilled at the bottom, were placed at opposite corners of the box. An unfamiliar adult male mouse was confined in one of the two beakers. The test started by placing the experimental mouse next to the beaker containing a male mouse and lasted 5 minutes. Distance moved and time spent at the beakers were analyzed using EthoVision. Rearing, self-grooming, and time spent sniffing the two beakers were analyzed using The Observer.

### 2.6 Accelerated Rotarod

Motor function was tested on a rotating rod with a diameter of 3.2 cm. The mice first underwent three pre-training trials, during which they were trained to remain on the rod at 3.75 rotations per minute (RPM) for up to 60 seconds. The mouse was placed on the rod during the training trials, which were performed at 3,75 RPM, accelerating to 40 RPM within 4 minutes. The maximum duration for each trial was 5 minutes. The test was interrupted if a mouse fell or could not balance on the rotating rod and grasp it, performing two consecutive rotations. The ITI was 45 minutes.

### 2.7 Pole Test

A vertical wooden stick (60 cm long, diameter 0.8 cm), stabilized on a wooden platform and marked by a black line every 20 cm, was used in the pole test. The markings divided the pole into three sections: 1 (upper level), 2 (middle level), and 3 (lower level). A mouse was placed with its head upward on the top of the pole. The nest was removed from its cage and placed on the platform to encourage the mice to climb. The latency of reaching the platform with all four paws defined the performance. Moreover, the mouse’s ability to turn 180° and climb down while keeping its head downward was evaluated. If the mouse turned, we recorded whether it turned at levels 1, 2, or 3.

### 2.8 Water maze

The Water maze test was used to assess the mice’s memory skills and consisted of one pre-training session and five training days. The pre-training was conducted under red light (5 lux): a 14-cm-diameter round platform was placed in the middle of a water-filled (21±1 °C) square pool (35 cm x 45 cm, 60 cm high), 1 cm below the water surface. During pre-training, the mice were familiarized with climbing onto the platform and waiting 10 seconds there before being picked up by the experimenter with a small shovel, thereby associating the platform with escape from the pool (Fellini et al., 2006). The training trials were conducted in a circular pool with a diameter of 145 cm. The water was kept at 21±1°C and colored white with non-toxic watercolor. The platform used during pre-training was placed 40 cm from the wall on the East side of the pool (Platform E or PE on Day 5; on the West side, Platform W or PW), 1 cm below the water surface. Four landmarks were placed behind the pool’s wall. Mice were gently placed into a non-transparent plastic cup, 5 cm in diameter, at the end of a 1 m-long plastic stick, and then placed into the pool from five positions in a pseudorandomized order. If a mouse did not find the platform within 90 seconds, the experimenter guided it to the platform, demonstrating that the mice became acquainted with the shovel during pre-training. After finding the platform and remaining there for 10 sec, the mice were presented with the shovel and climbed onto it, then returned to their home cages. During the recall trials, the platform was removed, the mice were placed in the center of the pool, and after 60 seconds, they were collected with a shovel at the previous platform position. An infrared light was projected onto the cages to keep the mice warm between trials. The experiments were video recorded and analyzed using EthoVision software. The protocol consisted of four experimental days: four training trials on day 1; two training trials and one recall trial (to assess short-term memory) on day 2; one recall trial (to assess long-term memory); and two training trials on day 3. Remote memory was evaluated using a recall trial on day 4 and 7 days after day 3.

### 2.9 Spontaneous intermale interactions

Social interactions between cagemate males were observed in their home cages at the animal facility. To distinguish the mice within a cage, each mouse’s tail was painted with an animal marking color. The behavior of the mice was observed by instantaneous sampling for 90 minutes with an interval of 3 min (thus, 30 samples/h) in the middle of the dark phase (14:00h). During the 3-minute intervals, the observer moved from one cage to the other and recorded the behavior displaced by each mouse in the cage. The observations were done on two days. On day one, the mice were placed in a new housing cage with fresh bedding. On day two, the mice were observed in a familiar cage where they had been housed for four days. The experimenter scored only parameters related to social interactions, namely: allogrooming (the mouse grooms a cage mate), social investigation (the mouse sniffs a cage mate), biting (the mouse bites a cage mate), fighting (the mouse chases a cage mate), mounting (the mouse mounts a cage mate), upright submissive posture (the mouse stands on the hind limbs showing the belly and neck to a cage mate).

### 2.10 DNA analysis and assessment of CAG repeats

DNA samples containing various CAG repeat lengths were generated by PCR from tail snips. First, DNA was purified (NucleoSpin gDNA Clean-up, Macherey-Nagel GmbH), and the concentration and purity of the DNA extracts were measured using a NanoDropTM 2000 (Thermo Fisher). Best PCR results were obtained under the following conditions: 100 ng genomic DNA, primers at a final concentration of 250 nM, 40 cycles of PCR, using AmpliTaq Gold™ 360 Mastermix (ThermoFisher), with 0,65 M betaine, 2.5% DMSO, and 50 µM deaza dGTP. The cycling conditions were 95°C 1 min, 5x touch down, 67°C, -1.5°C/cycle, followed by 35 x 95°C denat 30‘‘, 58°C annealing, 30‘‘, 72°C elongation 60‘‘. The primers used were 5’-6-Fam-GGC TGA GGA AGC TGA GGA -3’ as forward primer and 5’-GGC TGA GGA AGC TGA GGA -3’ as reverse primer, modified from Blanco et al. (2008). The resulting PCR products from the transgenic mice have 85-bp and 46-bp sequences flanking the CAG repeat core. Fluorescently labeled products obtained from the PCR reactions were analyzed on a 3130 Genetic Analyzer (Applied Biosystems) using POP-7 polymer. Electrophoresis was conducted with a GS-Matrix-G as a matrix file. Before the analysis, 2-4 µl of amplified product was added to 16-18 µl of deionized formamide containing 0.5 µl of GeneScan™ 600 LIZ™ Size Standard v2.0 as internal size standard with a 50 cm array. Samples were subjected to denaturation at 95°C for 3 min and then cooled on ice for 2 min before being loaded onto the Genetic Analyzer. The GeneScan profiles of the PCR products revealed a range of sizes; for our analyses, only the peak sizes were used. As has been described before (Williams et al., 1999; Duzdevich et al., 2011), the CAG repeat number measured by GeneMapper or PeakScanner differs from that measured by sequencing. Therefore, for the CAG repeat numbers determined by PeakScanner the following formula was applied (Duzdevich et al. 2011): True CAG repeat number = 1.0425 x PeakScanner CAG repeat number +1.2088. CAG sizes were calculated by subtracting the base pair length of the regions flanking the CAG repeat from the overall amplicon size as determined by Peak Scanner 2, converting to CAG units by dividing the corrected bp length by 3, multiplying the resulting CAG length by 1.045, and adding 1.2088, as described (Coffey et al., 2020).

### 2.11 Statistics

We performed two types of statistical analyses on all data. First, we analyzed the effects of genotype within each sex using the non-parametric Mann-Whitney test and, for repeated measurements, mixed 2-way ANOVA (with genotype as the between-groups factor and time bin, trial, day, or age as the within-groups factor). To test the possible effects of the interaction between genotype and sex, we performed 2-way ANOVA (with genotype and sex as between-groups factors) or mixed three-way ANOVA. For multifactor ANOVAs, Duncan post hoc comparisons were used when appropriate. One-sample tests against a given value (i.e., chance level for alternations in the Y-maze or the preference indexes in the social reference and spatial object recognition tests) were performed with the non-parametric Wilcoxon test. Correlation analyses were performed using the nonparametric Spearman test. All tests were two-tailed, and the significance level was set at 0.05.

## Results

### 3.1 Body weight is reduced in KI mice starting at 8 weeks of age

Body weight was reduced in the KI mice compared to the WT littermates (Fig. 1B for the raw data and Fig. 1C for data normalized to the average of the sex- and age-matched WT group). A mixed 2-way ANOVA detected a significant effect of genotype for both the female (F1,29= 17.910, P= 0.0002) and male (F1,26= 7.861, P< 0.0001) mice with no result of the interaction between genotype and age (females: F2,58= 0.8316, P= 0.595; males F2,52= 0.231, P= 0.795), indicating that the effect of the genotype did not depend on the age of the mice, as confirmed by Mann-Whitney comparisons between genotypes performed within each age group. A mixed 3-way ANOVA was performed on values normalized to WT mice within each sex and age group to control for consistent variability between male and female mice and age groups. There was a significant effect of genotype (F1,50= 21.52, P< 0.0001) and no effect of the interaction between genotype and sex (F1,50= 1.98; P= 0.171) and of the interaction between genotype, sex, and age (F2,100= 2.21; P= 0.115). Thus, KI mice weighed less than WT mice regardless of age and sex.

### 3.2 Novelty-induced exploration, motor coordination, social behavior, working, and short-term memory are not altered, while anxiety-related behavior is enhanced in 8-10-week-old KI mice

Novelty-induced behavior appeared to be not changed in 8-10 week-old KI mice as indicated by the parameters considered to be related to locomotionsuch as distance moved in the open field (Fig. 2A,B), time per transition in the Y-maze (Fig. 2F), total transitions in the EPM (Fig. 2G) and related to novelty-induced exploration such as rearing in the open field (Fig. 2E) and elevated plus maze (Fig. 2H), time at beakers of the social preference test (Fig. 2I), and time at objects of the spatial object recognition test (Fig. 2J,K). KI mice spent less time than WT mice on the open arms of the elevated plus maze (effect of genotype F1,52= 5.418; P= 0.024; Fig. 3A), and the impact of genotype seemed to be equal in male and female mice (effect of the interaction between genotype and sex F1,52= 0,292; P= 0.591). Notably, the percentage of open-arm entries did not differ between genotypes (Fig. 3B), indicating that KI mice visited the open arms as often as WT mice but did not explore them as extensively. Moreover, we detected a significant effect of the interaction between genotype and sex on the number of stretched attended postures (SAPs), a behavior indicative of risk asssesment towards an unfamiliar environment or stimulus, ( F1,52= 5.745; P= 0.020) (Fig. 3C) and time spent in the center (F1,52= 4.268; P= 0.044) (Fig. 3D): post-hoc analyses indicated that KI female mice more often displayed SAPs and spent more time in the center than WT mice indicating higher risk assessment towards the open arms. There was a significant effect of the interaction between genotype and sex (F1,52= 5.383; P= 0.024) on the percentage of full visits on the open arms (i.e., how often mice explored the total length of the arm reaching its end) (Fig. 3E) and on the latency to reach the end of one open arm (Fig 3F); posthoc comparisons indicated that KI female mice did less full visits and took longer time to reach the end of the open arms than WT female mice. Taken togerher, the results of the elevated plus maze suggest that female KI mice exhibited an enhanced risk assessment and reduced exploration of the open arms, which could be interpreted as increased anxiety compared to WT mice. On the other hand, we did not detect any differences in anxiety-related parameters in the open field, such as thigmotactic behavior (Fig. 2C) or at the border of the arena (Fig. 2D).

**Figure 2.**
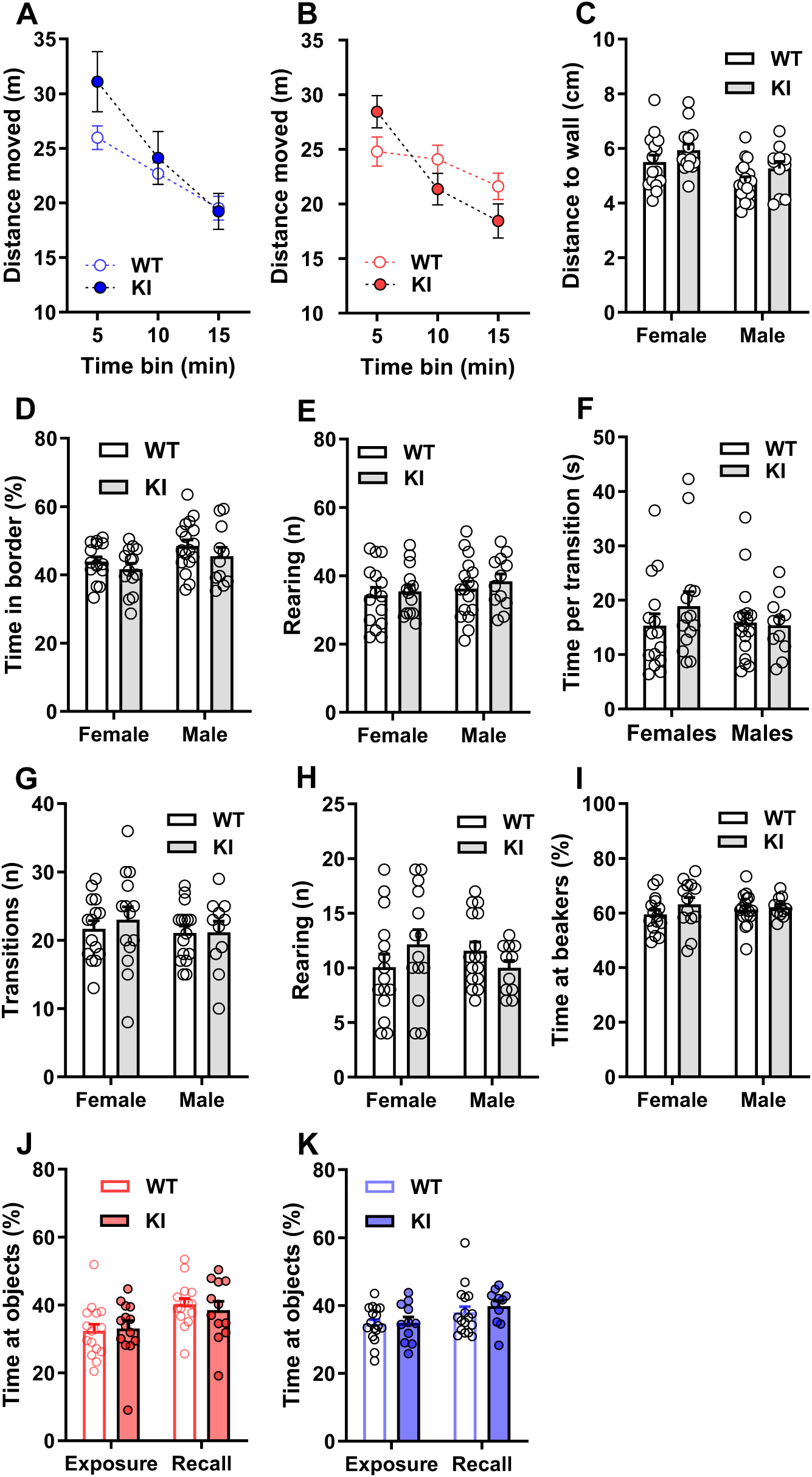
Unaltered novelty-induced exploration and locomotion in 8-to 10-week-old KI mice. A,B No difference was detected between WT and KI female (A) and male (B) mice in the distance moved in the open field test. C,D thigmotactic behavior was not affected by genotype as measured with the parameters mean minimal distance to the wall (C) and time spent in the border (D) of the open field. E, rearing during the first 5 minutes of the open field test was unaffected by genotype. F-K, novelty-induced locomotion and exploration were not affected by the genotype as indicated by the average time to make a transition in Y-maze of the spontaneous alternation test (F), number of transitions (G) and rearing (H) in the elevated plus maze, time spent at the beakers in the social interaction test (I), and time at the objects during the exposure and recall trials of the spatial object recognition test (J, female mice; K, male mice).

**Figure 3.**
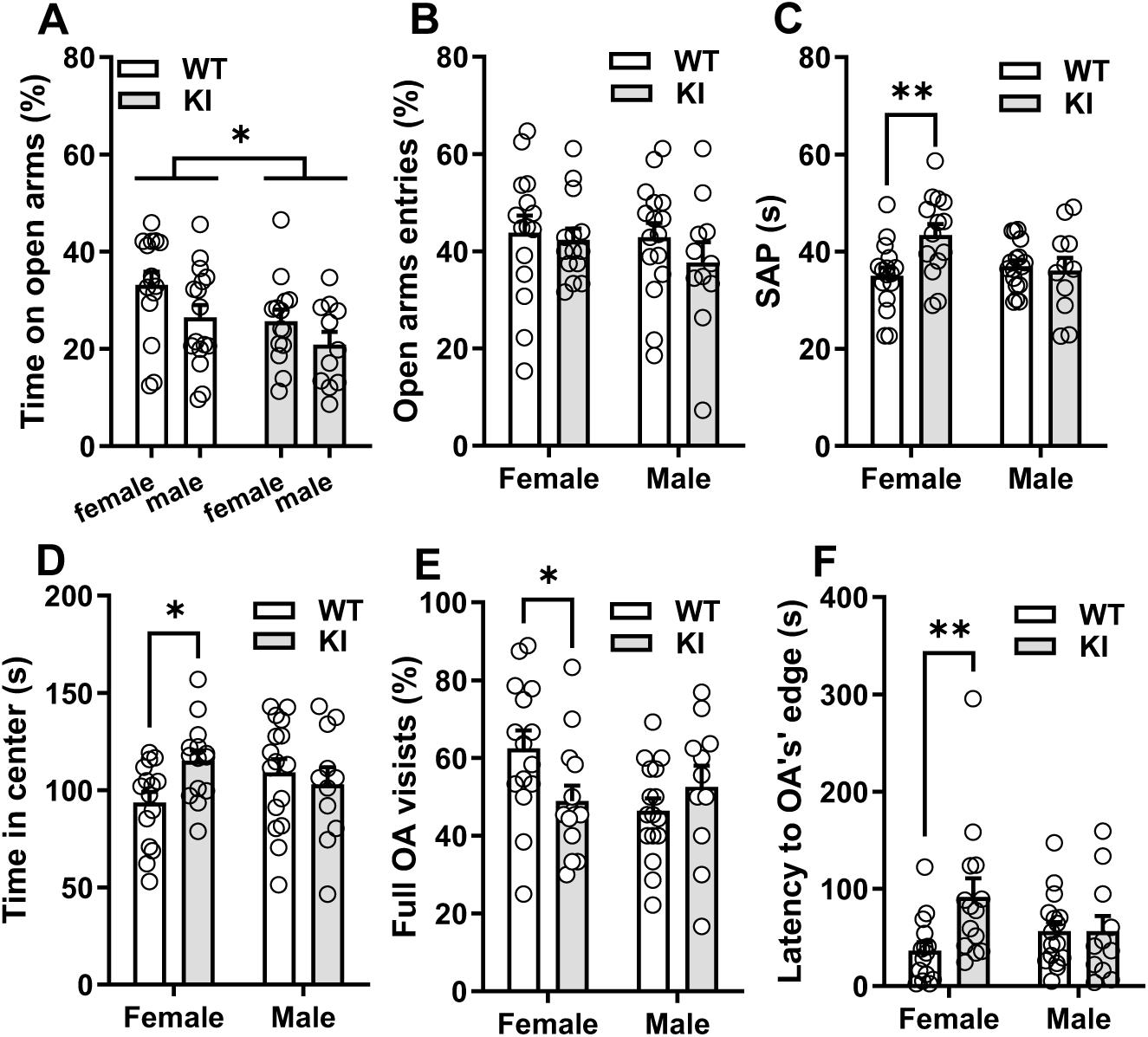
Enhanced risk assessment in the elevated plus maze in 9-week-old KI female mice. A, KI mice, regardless of sex, spent less time on the open arm of the elevated plus maze than WT mice (* p< 0.05, effect of genotype after the 2-way ANOVA). B, No difference between genotypes in the percentage of open arm entries. C, KI female mice spent a longer time in stretch attend posture (SAP) from the center towards the open arms than WT female mice (** p< 0.01, post-hoc test after the 2-way ANOVA). D, KI female mice spent a longer time in the center of the maze than WT female mice (* p< 0.05, post-hoc test after the 2-way ANOVA). E, KI female mice did not explore the entire length of the open arms as often as the WT female mice (* p< 0.05, posthoc test after the 2-way ANOVA). F, KI female mice took longer time than the WT female mice to enter for the first time one of the open arms (** p< 0.01, post-hoc test after the 2-way ANOVA).

In the social preference test, WT and KI mice spent more time at the beaker containing an unfamiliar sex-matched mouse than in the empty beaker, suggesting that social behavior was unaffected (Fig. 4A). Cognitive functions such as working memory and spatial short-term memory were normal in the KI mice, as indicated by the alternations in the Y-maze and the time spent at the displaced object in the spatial object recognition test, respectively (Fig. 4B-D). Finally, no differences between genotypes were observed in motor function and coordination, as assessed by the pole test and accelerated rotarod tests (Fig. 5A-E). The maximal velocity in the open field, which could also be used as an index of motor ability, did not differ between genotypes (Fig. 5F). Thus, novelty-induced exploration, social behavior, motor coordination, and cognitive functions such as working and short-term spatial memory appear unaltered in CAG140 KI mice during early adulthood.

**Figure 4.**
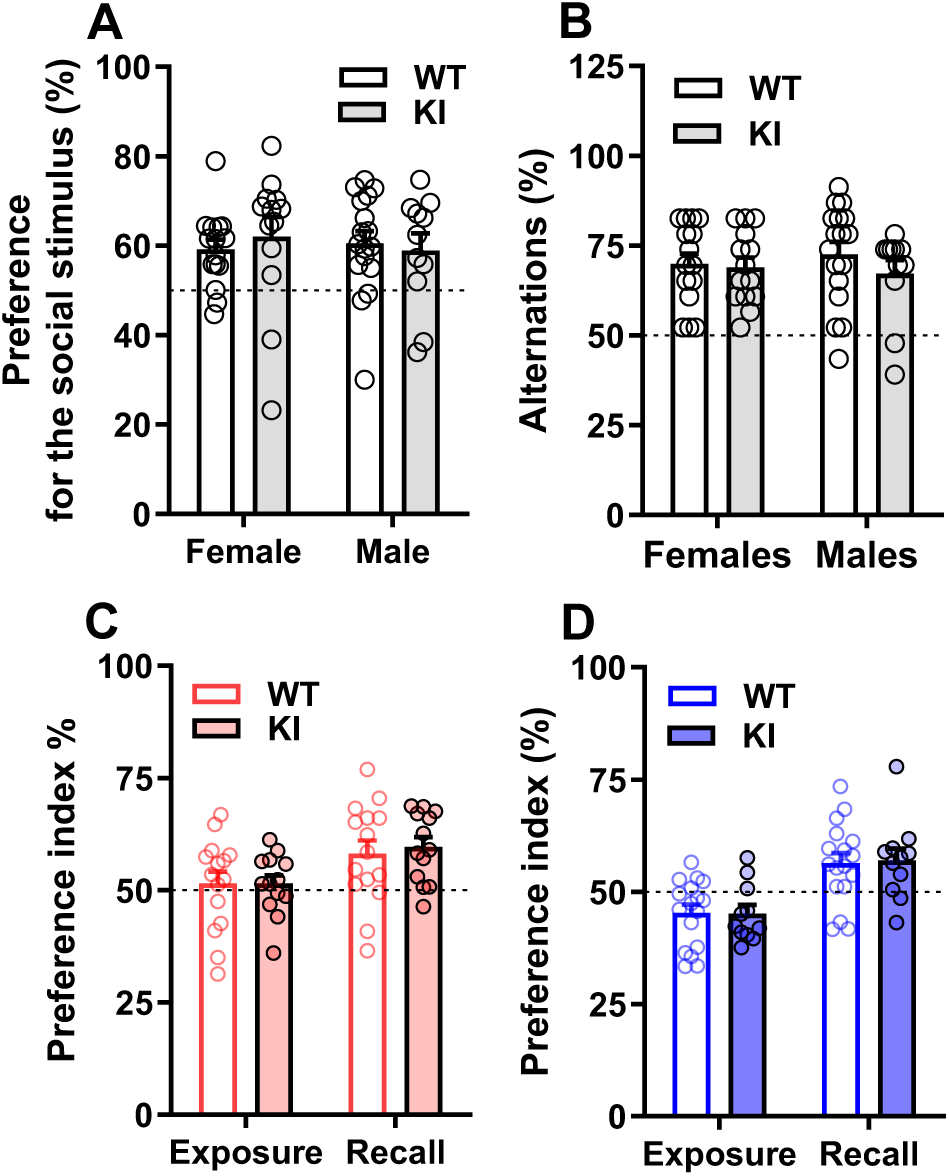
Social preference, working, and short-term memory are unaltered in 9-10-week-old KI mice. A, WT and KI mice spent more time investigating an unfamiliar sex-matched mouse than a novel object in the social preference test. B, working memory was unaffected in KI mice as tested in the spontaneous alternation test. C (female), F (male) WT, and KI mice preferred the spatially displaced object during the recall trial of the spatial object recognition tests.

**Figure 5.**
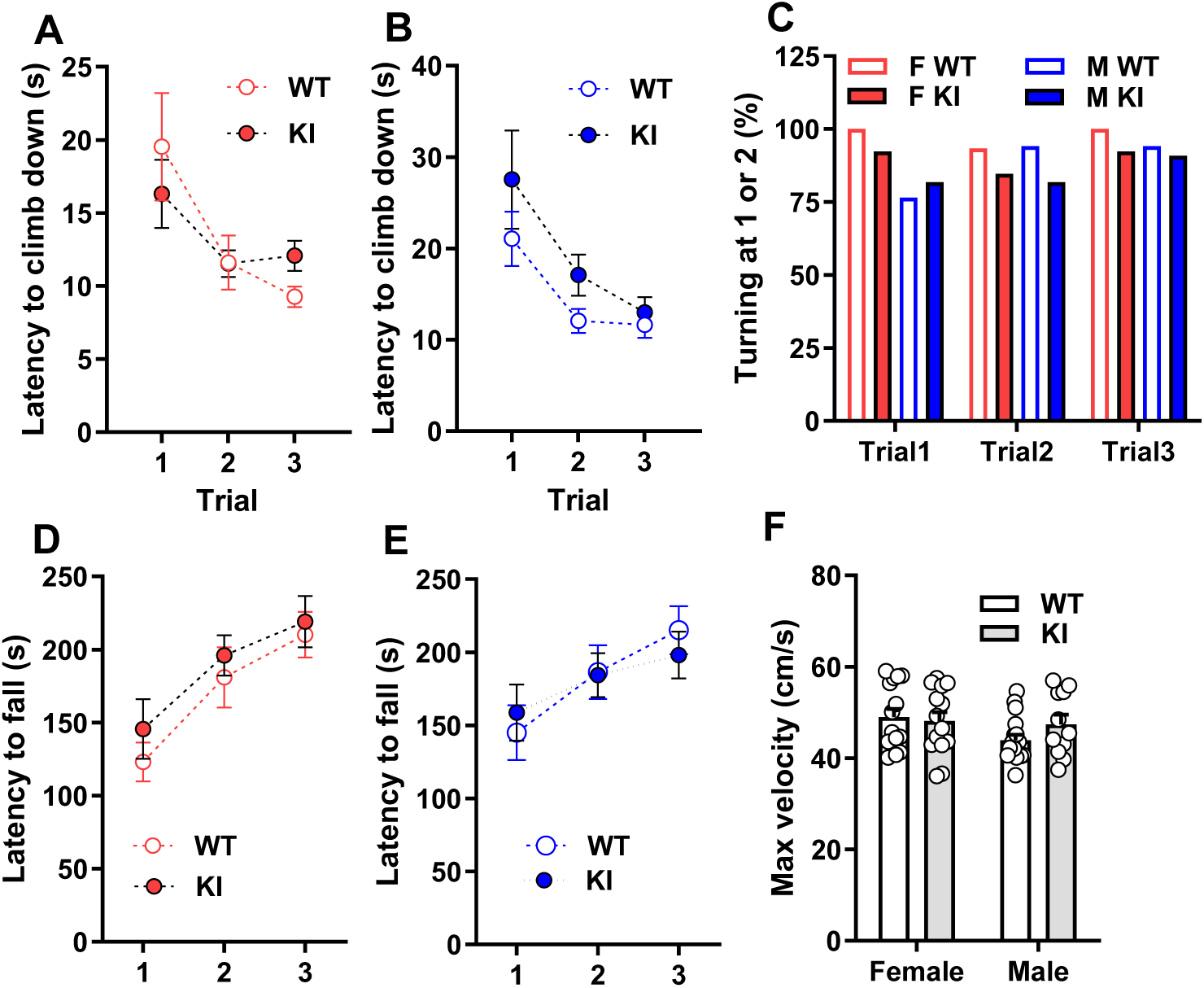
Motor function is not affected in 8- to 10-month-old KI mice. A, B, WT, and KI female (A) and male (B) mice showed shorter latencies to climb down from the first to the third trial of the pole test. C, most WT and KI mice could turn the body 180 degrees at the top of the pole before climbing down. D, E: female (D) and male (E) WT and KI increased latency to fall from the first to the third trial of the accelerated rotarod test. F, No differences between genotypes in their maximal velocity during the open field test performed at the age of 8 weeks.

### 3.3 Reduced locomotion in 14-16-week-old KI mice

When we tested the mice again at 14- to 16 weeks, we detected several significant differences between the genotypes (Fig. 6). KI mice moved less than WT mice in the open-field test, as indicated by a significant effect of genotype for both the female (F1,26= 4.848; P= 0.037) and male (F1,26= 4.754; P= 0.038) mice (Fig. 6A,B). Notably, there was no effect of the interaction between genotype and time bins (female F2,52= 0.320; P= 0.727; males F2,52= 0.830; P= 0.442), indicating that KI moved less than WT but showed similar short-term habituation to the arena. A mixed three-way ANOVA confirmed the effect of genotype (F1,53= 4.496; P= 0.039), the effect of the interaction between genotype and time bins (F2,106= 3.665; P= 0.029), and did not show any significant effect of the interaction between genotype and sex (F1,53= 1.621; P= 0.192). The maximal velocity in the open field was also reduced in KI mice compared to WT mice (effect of genotype: F1,52= 6.693; P= 0.012) independently of the sex of the mice (effect of the interaction between genotype and sex: F1,52= 0.024; P= 0.873) (Fig. 6C). No differences were detected for rearing, time spent in the center, and in the border (Fig. 6C-E). In line with the reduced locomotion in the open field, KI mice required, on average, more time to do a transition in the Y-maze compared to WT mice (effect of genotype: F1,52= 6.375; P= 0.015) independently of the sex (effect of the interaction between genotype and sex: F1,52= 0.460; P= 0.501) (Fig. 7A). In the Y-maze, both genotypes alternated at a rate above chance with no difference between genotypes, indicating that the transgenic CAG repeats did not affect working memory under the conditions of the spontaneous alternation task (Fig. 7B). In the pole test and accelerated rotarod, we did not detect any effect of genotype on the performance of the mice (Fig. 7C-D). When interactions within male cagemates at the age of about 16 weeks were observed, no differences were detected between genotypes, with both genotypes equally engaged in social and aggressive behaviors (Fig. 8). Together with the absence of altered social preference in 9-month-old mice, our data suggest that social and aggressive behaviors are not affected in the CAG140 line, indicating that this model does not capture the enhanced aggressiveness observed in Huntington’s disease patients (Fisher et al., 2014).

**Figure 6.**
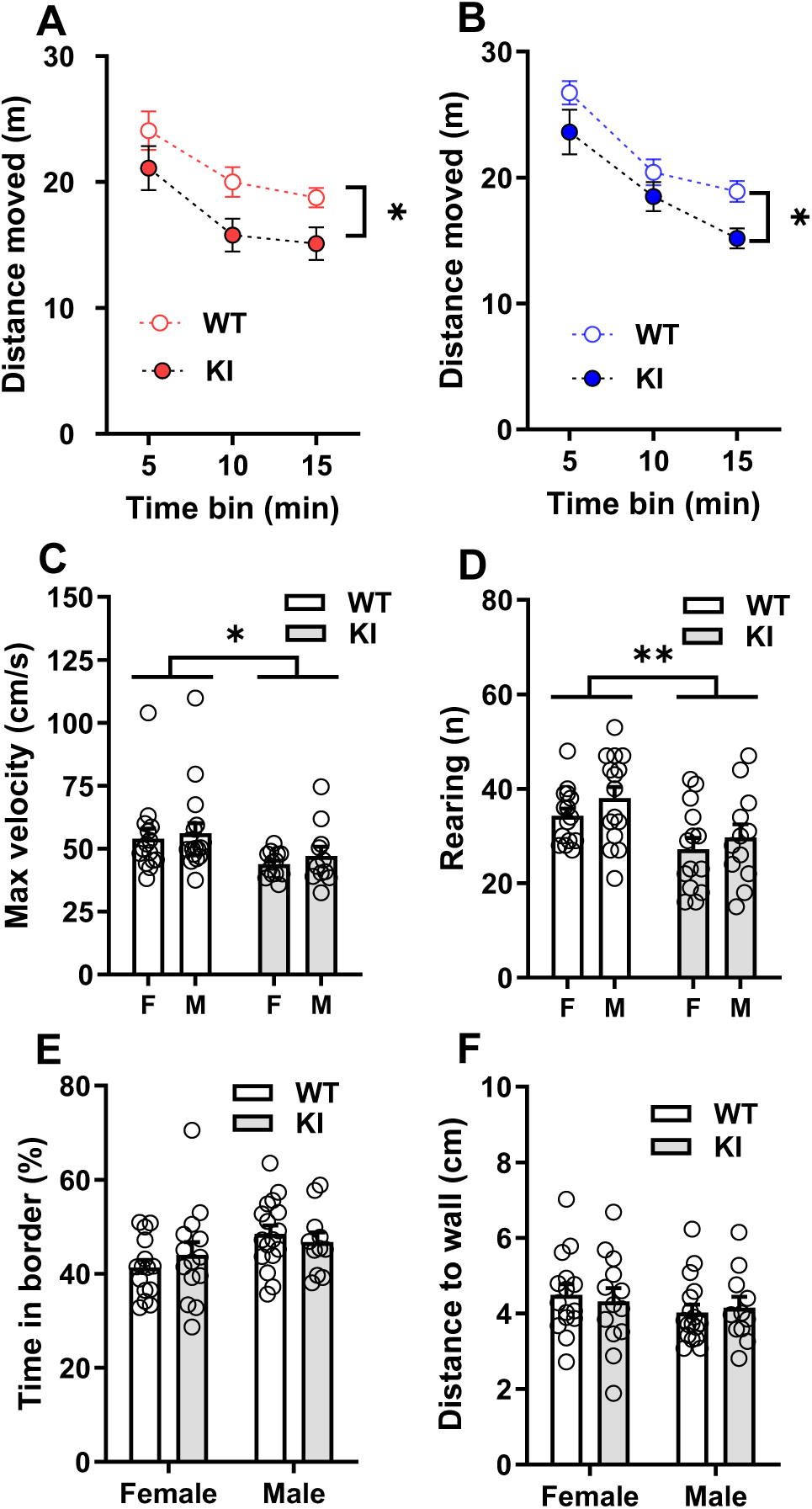
Decreased locomotion in the open field test in 14-week-old KI mice. A,B, female (A), and male (B) KI mice covered shorter distances in the open field test. C, the maximal velocity during the open field test was reduced in KI mice more than in WT mice. D, the KI mice did fewer rearing bouts in the open field than the WT mice. E,F, Thigmotactic behavior was not affected in KI mice, as shown by the time spent at the border (E) and the mean distance to the wall (F). * p< 0.05, ** p< 0.01 (effect of genotype after a 2- or mixed 3-way ANOVA).

**Figure 7.**
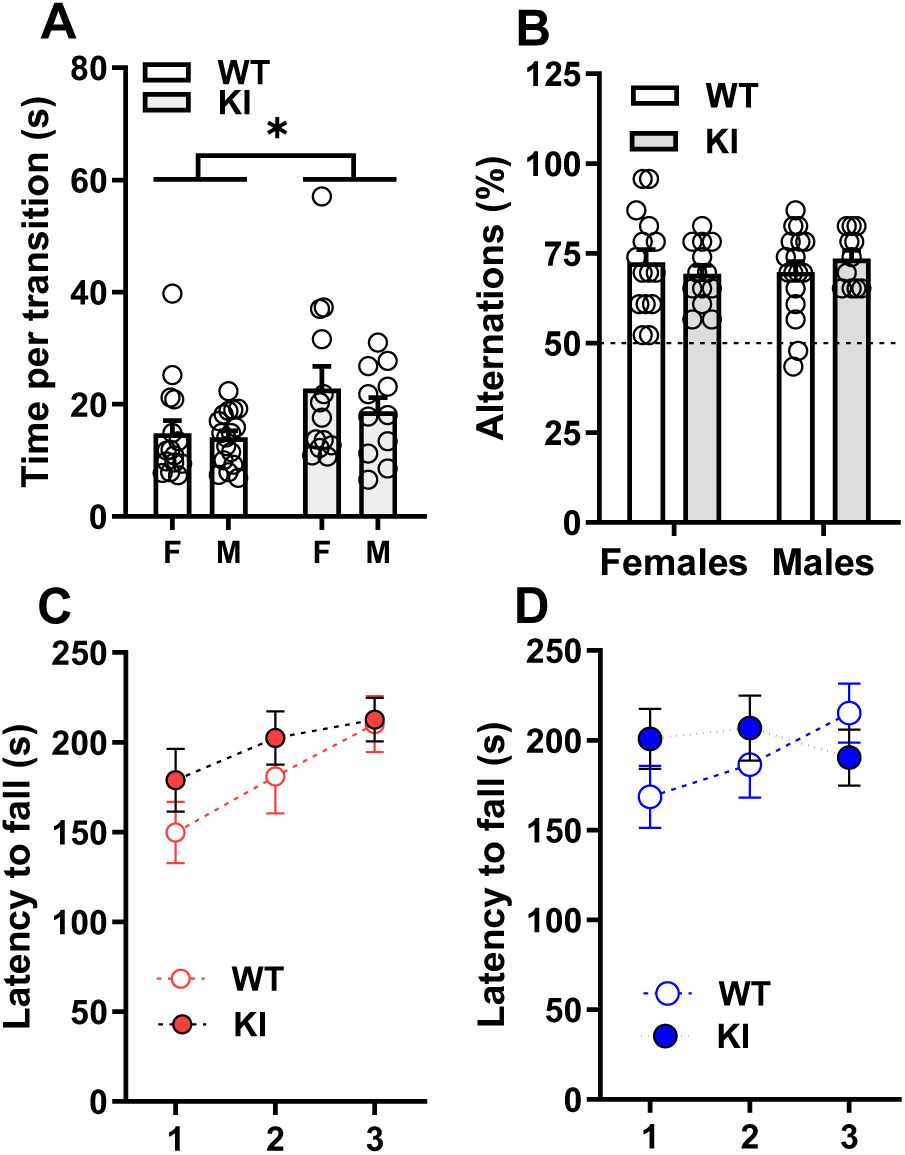
Unaltered working memory in the spontaneous alternation test and motor function in the rotarod test in the KI mice at the age of 15 weeks. A,B, in the spontaneous alternation test, KI mice show a similar percentage of alterations as the WT mice (B) but required more time to make a transition (A), indicating they were moving more slowly than WT mice. C, D, the latency to fall was not affected by the genotype within the female (C) and male (D) groups in the accelerated rotarod test. * p< 0.05 (effect of genotype after a 2-way ANOVA).

**Figure 8.**
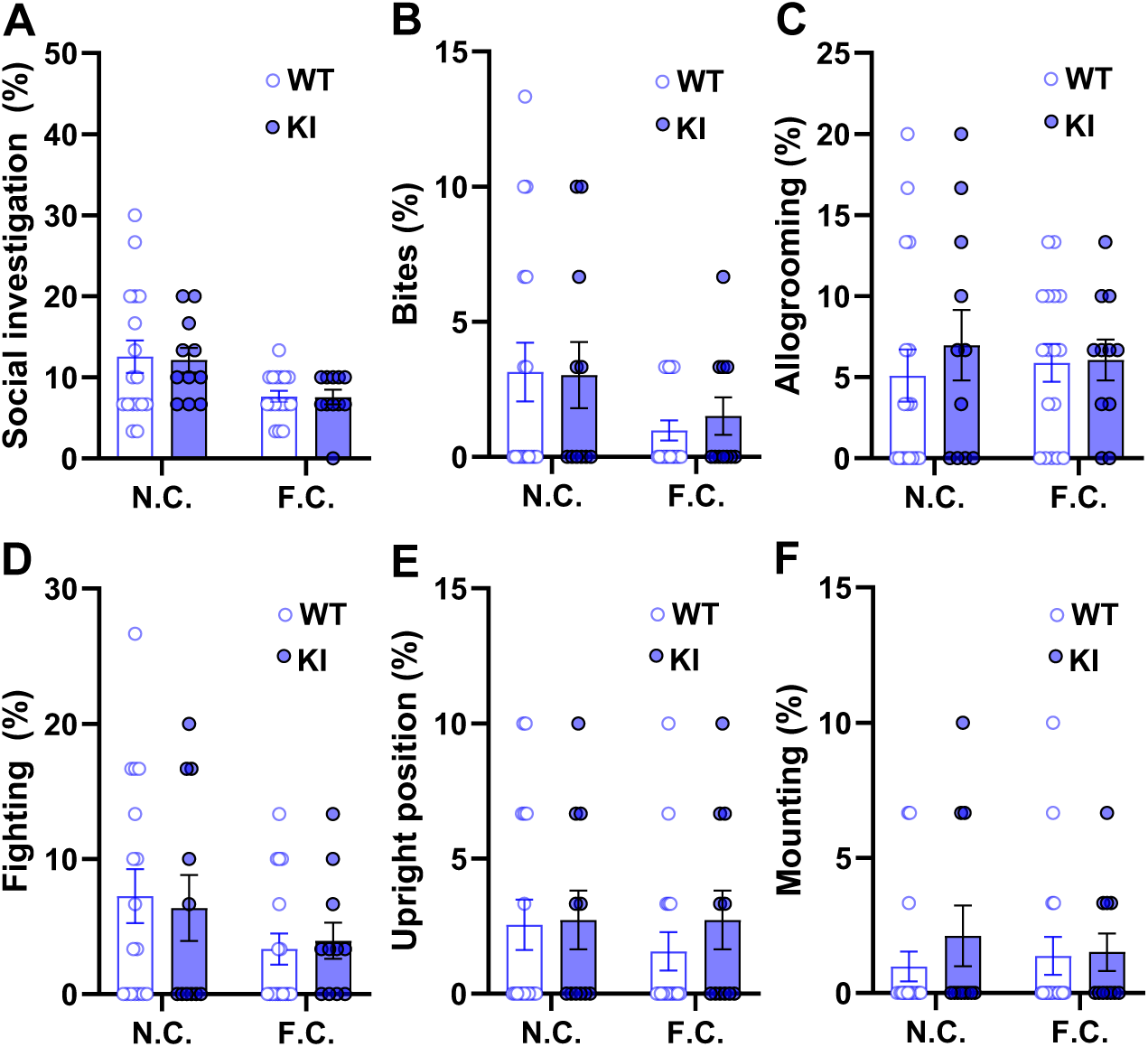
**Normal inter-males social and aggressive behavior in 16-week-old KI mice**. The spontaneous behavior of the mice in the home cage was analyzed 1 hour (new cage, N.C.) and 48 hours (familiar cage, F.C.) after they had been placed in a standard housing cage with fresh bedding. A-F, No differences between genotypes were detected in any social and aggressive behaviors analyzed.

### 3.4 Impaired long-term memory in 16-week-old KI mice

The water maze test was performed on 16-week-old mice to analyze their ability to acquire and retrieve a spatial map of the environment, a cognitive task considered to require the hippocampus and to model episodic memory in humans (for review, see Morellini, 2013). During the learning trials, WT and KI mice quickly improved their performance during the first training day and reached an asymptotic level at the end of the second training day (Fig. 9). A mixed 3-way ANOVA indicated no effects of the interaction between trial and genotype and of the interaction between trail, genotype and, sex on distance swum to reach the platform (genotype x trial F8,408= 0.544, P= 0.823; genotype x sex x trial F8,408= 0.690, P= 0.700; Fig. 9A,B). The mice underwent a short-term transfer trial 15 minutes after the second training trial on day 2 to assess the spatial strategies used to reach the escape platform. All groups preferred the target quadrant, spending more time than the chance level of 25%. No differences were detected between KI and WT female mice for all parameters indicative of a searching strategy in the proximity of the platform, namely time spent in the target quadrant (Fig. 9C), and mean minimal distance to the platform position (Fig. 9D). Thus, the analysis of the learning and transfer trials indicates that the genotype had no impact on the mice’s ability to learn the platform’s position. We performed two further transfer trials, 24 hours and 7 days after the last training trial, to test long-term and remote spatial memory. During these probe trials, we detected several differences between KI and WT mice within both sexes. Consistently, a three-way ANOVA detected a significant effect of the interaction between genotype and trials (i.e., for short-term, long-term, and remote memory) for two parameters indicative of a spatial preference for the platform position: time in the target quadrant (F2,104= 4.600, P= 0.012; Fig. 9B) and mean minimal distance to the platform position (effect genotype and trial F2,104= 3.320, P= 0.040; Fig. 9C), but no significant effect of the interaction between genotype and sex and trial for any of the analyzed parameters. The post hoc analyses indicated that, regardless of genotype, KI mice spent less time in the target quadrant and swam farther from the platform than WT mice during the probe trials for long-term and remote memory. Moreover, post hoc comparisons within genotypes indicated that KI spent less time in the target quadrant during long-term memory than during short-term memory (P= 0.049). Consequently, these findings suggest that although spatial learning and allothetic navigation remain intact in CAG140 KI mice, there is a specific impairment in long-term memory consolidation or retrieval.

**Figure 9.**
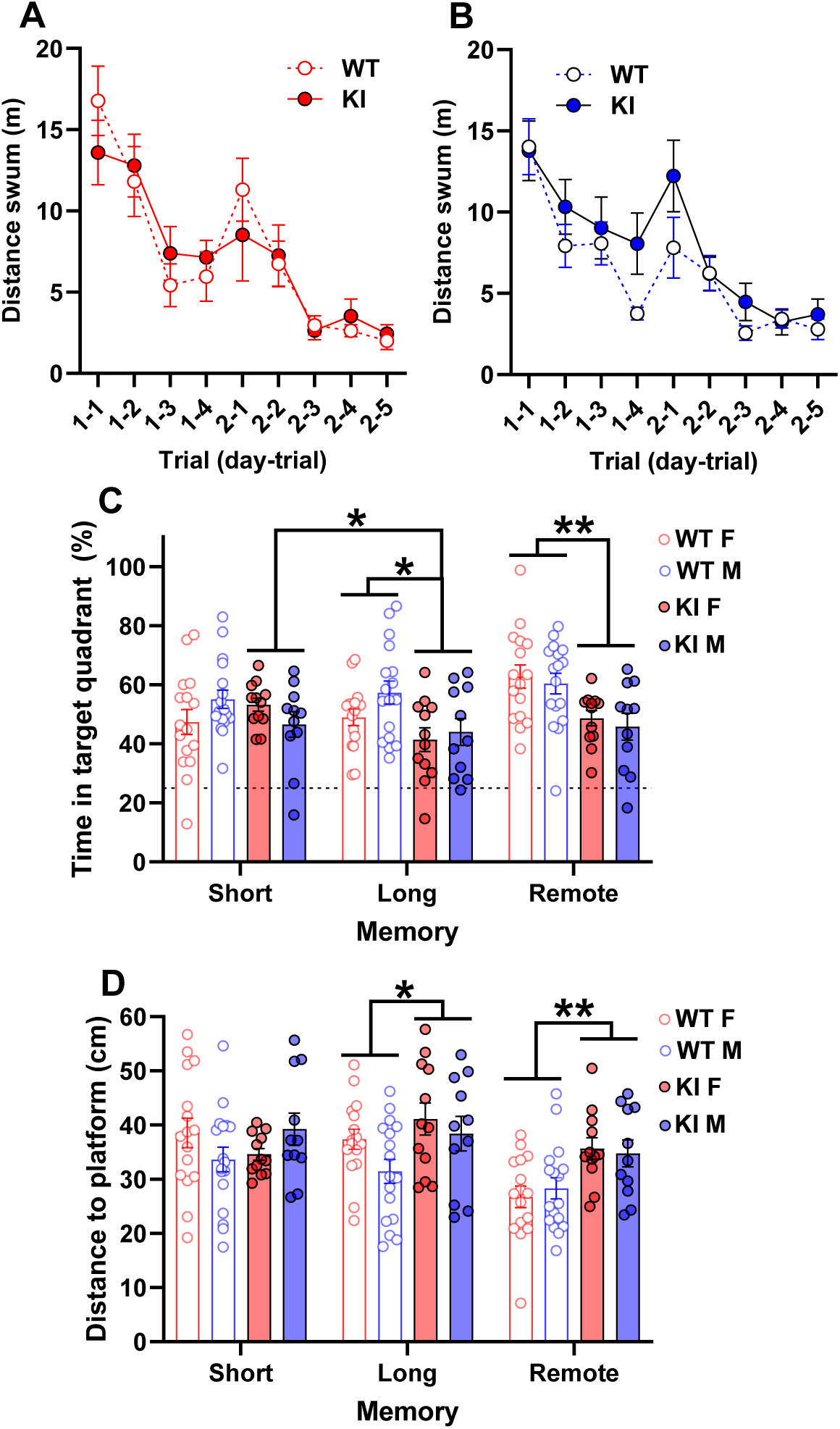
Impaired long and remote spatial memory in the KI mice at the age of 15 weeks. A,B, no difference during the learning trials between genotypes within the female (A) and male (B) sexes, as indicated by the distance swum to find the hidden platform in the water maze test. C and D, the mice underwent three probe trials at 20 min (short-term memory), 24 hours (long-term memory), and 14 days (remote memory) after the last training trial. All mice preferred the target quadrant, spending more time there than at chance level (25%, indicated by a dotted line). During the long-term and remote memory probe trials, the KI mice spent less time in the target quadrant (C) and searched at a longer distance from the platform (D) than the WT mice. Moreover, the KI mice reduced the time in the target quadrant from the short- to the long-term probe trial (C). * p< 0.05, ** p< 0.01 (post hoc comparisons after a significant effect of the interaction between trial and genotype calculated with a mixed 3-way ANOVA).

### 3.5 The degree of behavioral impairments correlates with the number of CAG repeats in the KI mice

Given the well-documented intergenerational instability of CAG repeats in the CAG140 lines, we aimed to investigate whether variations in repeat length exist within our experimental cohort of CAG140 mice; furthermore, it was of great interest to determine if these differences correlate with their behavior impairments, as consistently observed in human subjects (Andrew et al., 1993; Langbehn et al., 2004; Paulsen et al., 2008; Tabrizi et al., 2009). We thus performed cross-correlation analyses between body weight at 14 and 19 weeks, distance moved in the open field at 14 weeks, and time spent in the target quadrant during remote memory in WT and KI mice. In KI mice, but not in WT mice, the distance moved positively correlated with time in the target quadrant and with platform position. The CAG repeats varied from a minimum of 155 to a maximum of 165, with no differences between female (157.6 +/- 1.16) and male (157.3 +/- 0.96) mice (t19= 0.22, P= 0.829). The correlation analysis showed significant negative correlations between CAG repeats, distance moved in the open field (Fig. 10 C), and time spent in the target quadrant in the water maze test (Fig. 10 D).

**Figure 10.**
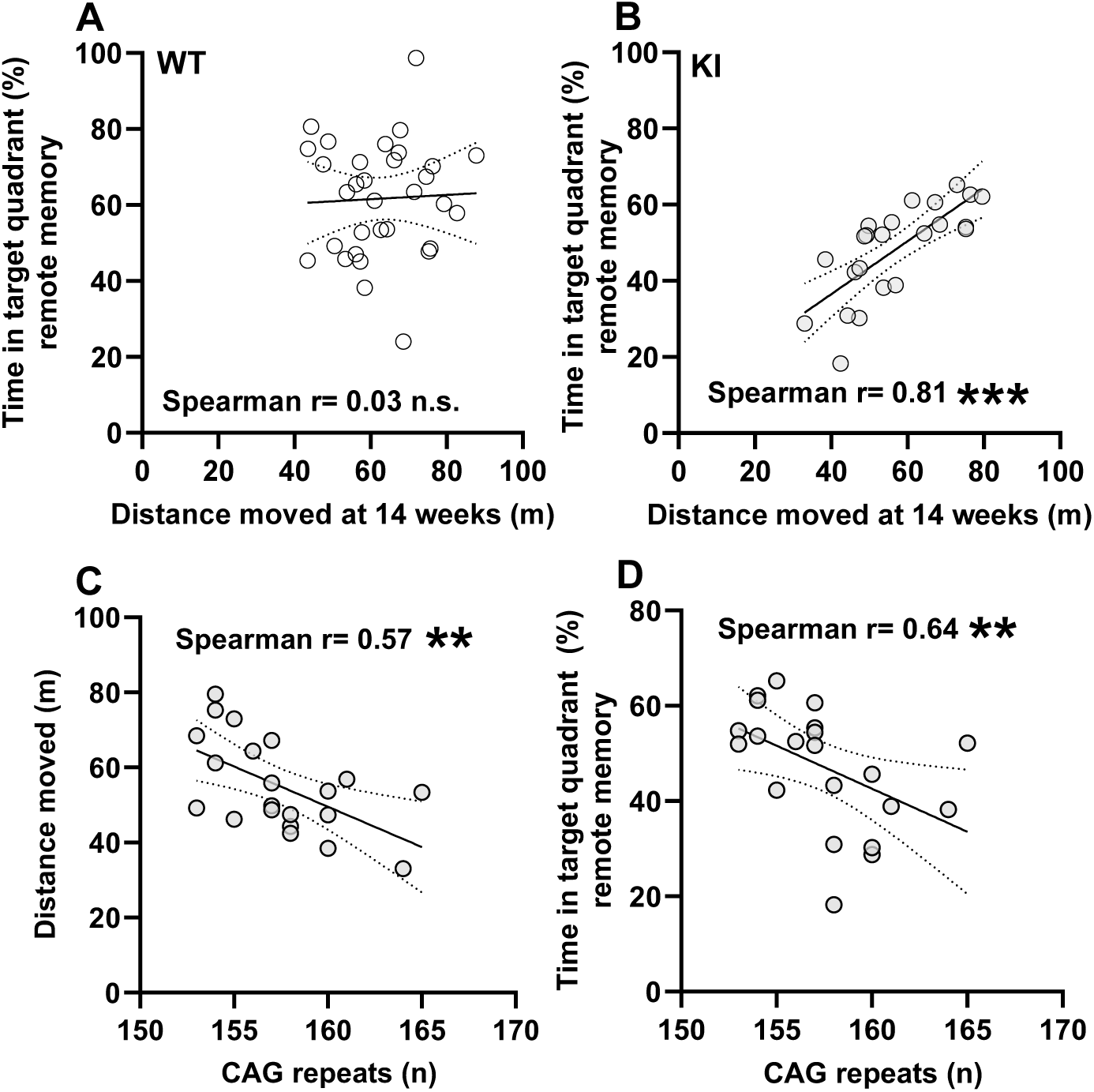
The number of CAG repeats correlates with behavioral alterations in KI mice. Remote memory correlates with locomotion and CAG repeats in the KI mice. A,B, distance moved in the open field positively correlates with the time spent in the target quadrant during the probe trial for remote memory of the water maze test in the KI (B) but not in the WT (A) mice. C,D, the number of the CAG repeats negatively correlates with the distance moved in the open field (C) and the time spent in the target quadrant during the probe trial of the water maze test (D). ** p< 0.01, *** p< 0.001 (Spearman correlation).

## Discussion

In the present study, we employed a validated mouse model of pathology in humans to test the hypothesis that preclinical symptoms, independent of the characteristic motor dysfunction in HD, can predict disease progression. Alterations in motor, emotional, and cognitive functions, as well as body weight, were identified in CAG140 KI mice, consistent with previous behavioral studies in this model of HD (Menalled et al., 2003; Hickey et al., 2008; Fowler and Muma, 2015; Stefanko et al., 2017; Rising et al., 2011). Notably, an intraindividual correlation was observed between phenotypic characteristics measured at different ages, with early-stage changes at 8 weeks predicting cognitive alterations at 16 weeks.

### Mild motor impairments in 14 weeks old CAG140 KI mice

Reduced locomotion and rearing behavior in CAG140 KI mice were associated with decreased maximal ambulatory velocity, suggesting minor motor deficits at 14 weeks of age. However, no impairments were detected in the pole and rotarod tests, which assess motor coordination. This dissociation between spontaneous locomotion and coordinated motor performance suggests that open field impairments may involve motivational or affective components rather than purely motor deficits. Supporting this interpretation, a slight but significant increase in anxiety-like behavior was observed on the elevated plus maze in 10-week-old CAG140 KI mice. While most studies on CAG140 mice have not specifically analyzed anxiety, Hickey et al. (2008) reported increased anxiety in the light/dark box and enhanced freezing during fear conditioning. Thus, heightened anxiety may underlie the reduced locomotion, maximal velocity, and rearing observed in the open field at 14 weeks, consistent with psychiatric symptoms frequently reported in HD, such as anxiety and apathy (Craufurd, Thompson, and Snowden, 2001; van Duijn, Kingma, and van der Mast, 2007).

The absence of impaired motor function in the rotarod and vertical pole tests observed in this study aligns with previous research, which reported motor coordination deficits at later ages than those investigated in our study, including gait abnormalities at one year (Menalled et al., 2013), impaired performance in the accelerated rotarod beginning at one year (Rising et al., 2011), and tremor-like in-place movements at 65 weeks (Fowler and Muma, 2015). In younger mice, motor impairments in CAG140 KI mice have primarily been inferred from reduced spontaneous locomotion and diminished rearing behavior (Menalled et al., 2003; Fowler and Muma, 2015; Stefanko et al., 2017; Rising et al., 2011), both of which were also observed in the present study. The only exception is the study by Hickey et al. (2008), which identified subtle rotarod impairments exclusively under specific protocols and with large sample sizes.

### CAG140 KI mice show a specific impairment in long-term memory at the age of 16 weeks

The relatively mild motor deficits observed in CAG140 mice in our study enabled us to investigate other HD-associated behavioral alterations without confounding effects of reduced motility. In tests of cognitive function, CAG140 KI mice exhibited normal short-term and working memory but significant impairments in long-term memory, indicating selective vulnerability of neural circuits involved in memory consolidation and retrieval. This finding represents the first evidence of impaired long-term memory in CAG140 KI mice. Previous studies assessing short- and long-term memory using novel object recognition did not report alterations in 4- and 6-month-old mice (Stefanko et al., 2017). This discrepancy may be due to the reliance of novel object recognition on familiarity, a cognitive process independent of the hippocampus, in contrast to the spatial version of the water maze used in the present study (Good et al., 2007; Albani et al., 2014; Morellini, 2013). Notably, in the same study, CAG140 KI mice at exhibited specific impairments in the reversal learning protocol of the Y-maze, while performing comparably to controls during the initial learning phase (Stefanko et al., 2017), further supporting the hypothesis of impaired hippocampal function that is known to play an important role in reversal learning (Watson et al., 2009).

The early deficits in long-term and episodic-like memory observed in CAG140 KI mice may be clinically relevant, as they mirror cognitive changes commonly reported in patients with HD, where such impairments often emerge during prodromal or early symptomatic stages (Hamilton et al., 2003; Tabrizi et al., 2009). However, the dissociation between preserved short-term and working memory and impaired long-term memory in CAG140 KI mice contrasts with the typical clinical presentation, in which working and short-term memory are also frequently compromised in presymptomatic individuals (Guillot et al., 2025; Martinez-Horta et al., 2025; Stout et al., 2011; Paulsen et al., 2008). These findings highlight both the translational value and the limitations of the CAG140 model, as it reflects only a subset of the cognitive deficits observed in human patients. Mechanistically, long-term memory deficits in CAG140 KI mice likely result from combined hippocampal and cortical dysfunction, including synaptic alterations and neurotransmitter imbalances. CAG140 KI mice exhibit hypoperfusion in the prelimbic and cingulate cortex, as well as the hippocampus, at 6 months of age, prior to the onset of motor deficits and in the absence of striatal atrophy (Wang et al., 2016). Collectively, these findings provide strong evidence that cortical and hippocampal dysfunction precedes motor changes, consistent with presymptomatic cognitive deficits in HD.

### Memory impairments in 16-month-old CAG140 KI mice are predicted by their motor behavior at the age of 8 month and correlate with CAG repeats length

A significant correlation between spontaneous locomotion and cognitive deficits was observed, indicating that the neurodegenerative processes underlying HD may simultaneously affect multiple functional domains. These findings align with the concept that HD pathology, including neuronal loss and dysfunction in key brain regions such as the striatum and cortex, leads to a spectrum of behavioral impairments encompassing motor, emotional, and cognitive functions (Hodges et al., 2006). Another key finding of this study is the correlation between behavioral deficits and CAG repeat length: mice with longer CAG tracts exhibited more pronounced impairments in both locomotion and memory, consistent with dose-dependent effects observed in other models (Mangiarini et al., 1996; Slow et al., 2003). This relationship supports the hypothesis that polyglutamine expansion exerts a dose-dependent effect on the severity of HD symptoms. Similar findings have been reported in other mouse models, where the extent of CAG repeat expansion correlates with the onset and severity of motor and cognitive dysfunction (Lin et al., 2001; Menalled et al., 2012). In humans, longer CAG repeats in the *HTT* gene are associated with earlier disease onset and increased disease burden, including progressive motor and cognitive impairments (Andrew et al., 1993; Langbehn et al., 2004; Paulsen et al., 2008; Tabrizi et al., 2009). These correlations highlight the genetic basis of disease variability and emphasize the importance of CAG repeat length as both a pathogenic factor and a potential biomarker for disease progression.

Notably, we detected CAG repeat numbers between 155 and 165, slightly elevated relative to the original reports for the CAG140 line. This discrepancy could be caused by the substantial intergenerational CAG repeat instability, with published measurements ranging from 122–147 CAGs in standard colonies (Fowler and Muma, 2015) and 130 ± 3 CAGs in independent reports (Corey-Bloom et al., 2017), and even 121 CAGs in early generations (Hockey et al., 2008). Spontaneous expansions have also been observed, giving rise to CAG140-derived lines such as zQ175 with 176 ± 8 to 190 repeats (Gray et al., 2024; Peng et al., 2016; Lim et al., 2020). Thus, our measured values (155–165 CAGs) fall within the documented variability of this genetic background and are consistent with known drift in CAG140 alleles. Given the robust correlations observed between CAG repeat length and behavioral severity, coupled with the marked intergenerational and inter-colony instability demonstrated in this animal model, routine internal validation of CAG repeat numbers represents a highly recommended practice for future investigations.

In summary, the CAG140 mouse model provides valuable insights into the behavioral and cognitive manifestations of HD. The combination of reduced locomotor velocity, mild anxiety, and selective long-term memory deficits reflects early clinical stages of HD and highlights the interdependence of motor, cognitive, and affective symptoms. The presence of subtle emotional and cognitive impairments in CAG140 mice at an age when major motor symptoms are absent aligns with reports of cognitive and behavioral symptoms emerging up to 15 years before onset of motor dysfunction in HD patients (for review, see Paulsen, 2013). Collectively, the data suggets that CAG140 mice are well suited to elucidate mechanisms underlying HD pathology and for testing targeted interventions to mitigate both motor and cognitive decline.

## 5. Acknowledgemets

The authors are grateful to Tanja Stoessner and Karen Kesseler for excellent mouse husbandry. This work was supported by grants from the Deutsche Forschungsgemeinschaft CRC1436 TPA02 and from the European Research Council (ERC) under the European Union’s Horizon Europe research and innovation programme (grant agreement No. 101161748, acronym LEXSYN) to KMG; CRC1436 TPA02, TPA04, Z01, FG 5228 Syntophagy RP6, DFG graduate program SynAge, Alzheimer Forschung Initiative (AFI Grant Number: #23055R), RGP0002/2022 HFSP to MRK.

## 6. Authors contrbtions

**Olga Jung:** Conceptualization, Investigation, Visualization, Writing-Original draft preparation. **Sabine Hoffmeister-Ullerich:** Investigation. **Asina Omriouate:** Investigation. **Josefine Plumhoff:** Investigation. **Katarzyna M. Grochowska:** Conceptualization, Funding acquisition. **Michael Kreutz:** Conceptualization, Funding acquisition. **Fabio Morellini:** Conceptualization, Supervision, Visualization, Writing-Original draft preparation. **All authors:** Writing-Reviewing and Editing.

## 7. Disclusure

Figure 1A was generated with the assistance of OpenAI ChatGPT image generation tools (GPT-5.5/DALL·E) based on author-defined specifications. The authors reviewed and modified the generated output to ensure scientific accuracy and accept full responsibility for the final figure.

